# *De novo* Gene Signature Identification from Single-Cell RNA-Seq with Hierarchical Poisson Factorization

**DOI:** 10.1101/367003

**Authors:** Hanna Mendes Levitin, Jinzhou Yuan, Yim Ling Cheng, Francisco J.R. Ruiz, Erin C. Bush, Jeffrey N. Bruce, Peter Canoll, Antonio Iavarone, Anna Lasorella, David M. Blei, Peter A. Sims

## Abstract

Common approaches to gene signature discovery in single cell RNA-sequencing (scRNA-seq) depend upon predefined structures like clusters or pseudo-temporal order, require prior normalization, or do not account for the sparsity of single cell data. We present single cell Hierarchical Poisson Factorization (scHPF), a Bayesian factorization method that adapts Hierarchical Poisson Factorization [1] for *de novo* discovery of both continuous and discrete expression patterns from scRNA-seq. scHPF does not require prior normalization and captures statistical properties of single cell data better than other methods in benchmark datasets. Applied to scRNA-seq of the core and margin of a high-grade glioma, scHPF uncovers marked differences in the abundance of glioma subpopulations across tumor regions and subtle, regionally-associated expression biases within glioma subpopulations. scHFP revealed an expression signature that was spatially biased towards the glioma-infiltrated margins and associated with inferior survival in glioblastoma.

## Introduction

Recent advances in the scalability of single cell RNA-sequencing (scRNA-seq) offer a new window into development, the cellular diversity of complex tissues, cellular response to stimuli, and human disease. Conventional methods for cell-type discovery find clusters of cells with similar expression profiles, followed by statistical analysis to identify subpopulation-specific markers [2-5]. Studies of cell fate specification have benefitted from innovative methods for inferring pseudo-temporal orderings of cells, allowing identification of genes that vary along a trajectory [6-8]. By design, these approaches discover expression programs associated with either discrete subpopulations or ordered phenotypes like differentiation status. However, in addition to cell type and developmental maturity, a cell’s transcriptional state may include physiological processes like metabolism, growth, stress, and cell cycle; widespread transcriptional alterations due to copy number variants; and other co-regulated genes not specific to a discrete subpopulation or temporal ordering. Such expression programs are of particular interest in diseased tissue, where the underlying population structure may be unknown and druggable targets might vary independently of cell type or maturity.

Matrix factorization is an appealing approach to decomposing the transcriptional programs that underlie cellular identity and state without a predefined structure across cells. In this class of models, both cells and genes are projected into the same lower-dimensional space, and gene expression from each cell is distributed across latent factors that approximate a vector basis for its transcriptional profile. Genes’ weights over the latent factors are discovered simultaneously and can be used to identify expression programs. For example, previous studies have defined gene expression programs from scRNA-seq data using Principal Component Analysis (PCA) or non-negative matrix factorization (NMF) [9-13]. However, a combination of biological regulation, stochastic gene expression, and incomplete experimental sampling leads to sparsity in scRNA-seq data. This creates challenges in downstream analysis. Conventional methods like PCA and NMF are sensitive to false-negative dropout events in which a transcript is experimentally undetected despite its presence in a cell [14, 15]. Further, sparsity may vary across both cells and genes, complicating the normalization that most computational methods require [15, 16].

Here, we describe single-cell Hierarchical Poisson Factorization (scHPF), a Bayesian factorization method that uses Hierarchical Poisson Factorization [1] to avoid prior normalization and explicitly model variable sparsity across both cells and genes. We compare scHPF to popular normalization and dimensionality reduction methods as well as an algorithm explicitly designed for scRNA-seq. scHPF has better predictive performance than these methods and more closely captures expression variability in datasets generated by multiple experimental technologies. Finally, we apply scHPF to single-cell expression profiles obtained from the core and invasive edge of a high-grade glioma. scHPF identifies both expected and novel features of tumor cells at single-cell resolution and uncovers a prognostic expression signature associated with poor survival in glioblastoma.

## Results

### Single-cell Hierarchical Poisson Factorization

scHPF uses Hierarchical Poisson Factorization [1] for *de novo* identification of gene expression programs. In scHPF, each cell or gene has a limited “budget” which it distributes across the latent factors (**Fig. 1**). In cells, this budget is constrained by transcriptional output and experimental sampling. Symmetrically, a gene’s budget reflects its sparsity due to overall expression level, sampling, and variable detection. The interaction of a given cell and gene’s budgeted loadings over factors determines the number of molecules of the gene detected in the cell.

**Figure 1:**
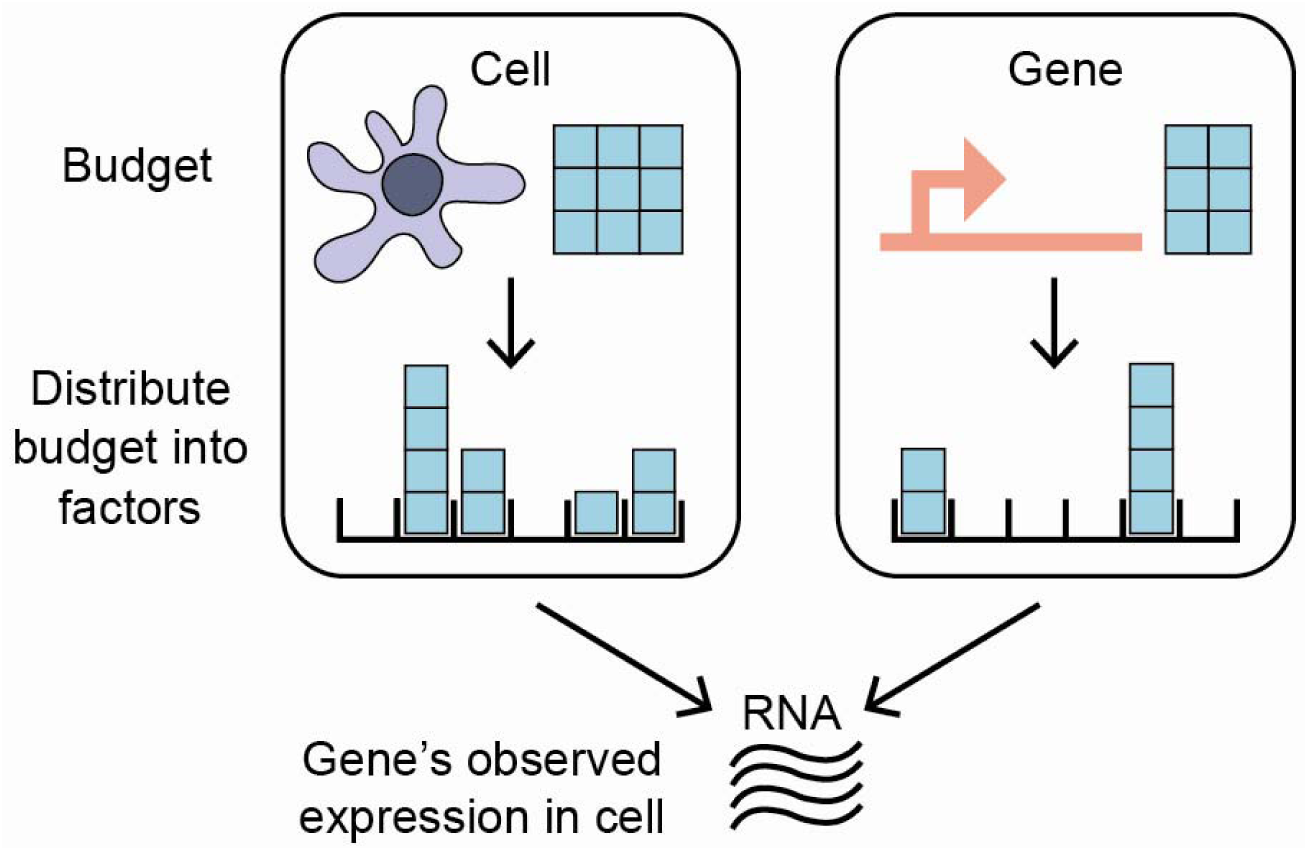
Cartoon representation of cells and genes allocating “budgets” across latent factors. The interaction of a cell and gene’s budget-constrained loadings over factors determines the gene’s observed expression level in the cell.

More formally, scHPF is a hierarchical Bayesian model of the generative process for an M x N discrete expression matrix, where M is the number of genes and N is the number of cells (**Fig. S1a**). scHPF assumes that each gene *g* and cell *c* is associated with an inverse-budget *η_g_* and *ξ_c_* that probabilistically determines the observed transcriptional output for that cell or gene. Since both *η_g_* and *ξ_c_* are positive-valued, scHPF places Gamma distributions over those latent variables. The hyperparameters of these Gamma distributions are set empirically (Methods, **Fig. S1b**). For each factor *k*, gene and cell loadings over factors *β_g,k_* and *θ_c,k_* are drawn from a second layer of Gamma distributions whose parameters depend on the inverse budgets *η_g_* and *ξ_c_* for each gene and cell. Finally, scHPF posits that the observed expression of a gene in a given cell is drawn from a Poisson distribution whose rate is the inner product of the gene’s and cell’s weights over factors. Importantly, scHPF accommodates the over-dispersion commonly associated with RNA-seq [17] because a Gamma-Poisson mixture distribution results in a negative binomial distribution; therefore, scHPF implicitly contains a negative binomial distribution in its generative process. Given a gene expression matrix, scHPF approximates the posterior distribution over the inverse budgets and latent factors given the data using Coordinate Ascent Variational Inference [18, 19] (Methods). After fitting the model’s variational posterior, we define each gene and cell’s score for a factor *k* as the expected values of its factor loading *β_g,k_* or *θ_c,k_* times its inverse budget *η_g_* or *ξ_c_*, respectively. We select the number of factors based on the convergence of the negative log likelihood and representation of major cell types (Methods).

### Benchmarking against alternative methods

We compared scHPF’s predictive performance to that of PCA, NMF, Factor Analysis (FA), and Zero Inflated Factor Analysis (ZIFA) [14], a method developed specifically for scRNA-seq. These methods have been used for *de novo* expression program discovery without a pre-defined structure across cells [9-12, 14]. We assessed each method across three datasets in different biological systems and obtained with different experimental platforms (**Table S1**). The peripheral blood mononuclear cell (PBMC) data from 10x Genomics is a mixture of discrete cell types [20], whereas the Matcovitch *et al*. microglial dataset samples from multiple timepoints along a developmental process [21]. Additionally, we profiled 9,924 cells from a patient-derived glioma neurosphere line (TS543), in which physiological processes like cell cycle, rather than discrete cell-types or differentiation status, drive expression variability. The datasets originate from different biological systems and experimental technologies including: droplet-based 10x Chromium [20], MARS-seq [22], and an automated microwell platform [23].

For each dataset, we tested conventional methods with three different normalizations: log-transformed molecular counts, counts per median (rate-normalization), and log-transformed counts per median (log-rate-normalization). ZIFA was only evaluated using log-transformed normalizations as recommended by its authors. Across all datasets and normalizations, scHPF had the best predictive performance on a held-out test set (**Fig. 2a**). scHPF’s superior performance was robust across a range of values for *K*, the number of factors (**Fig. S2**).

**Figure 2:**
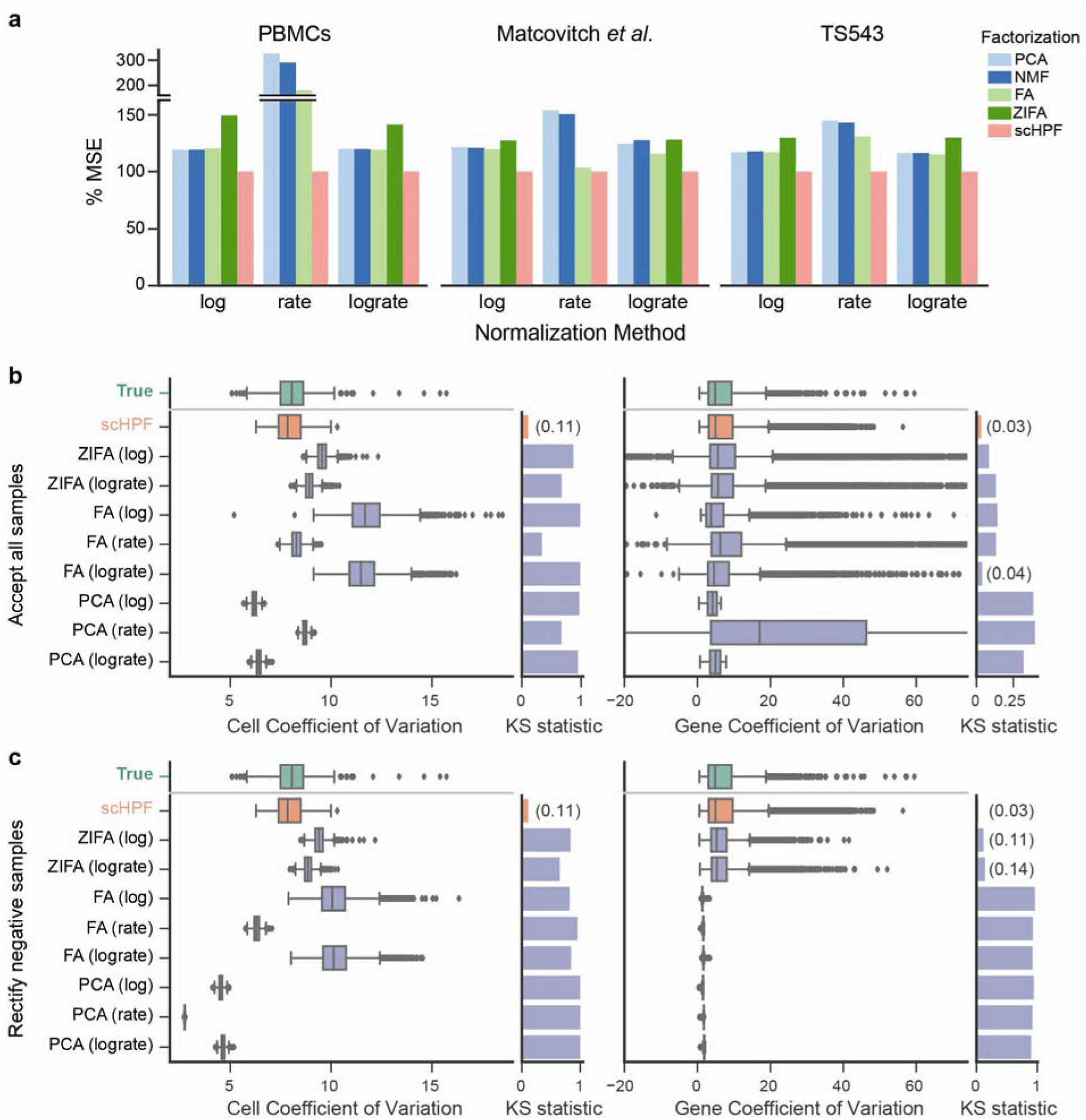
scHPF captures statistical properties of scRNA-seq data better than alternative factorization methods. **(a)** Mean squared error (MSE) of different factorization methods on a withheld test set as a percent of scHPF’s. scHPF’s MSE was calculated after normalizing its predictions. **(b)** Posterior predictive checks of expression variability in PBMCs. Box plots show the coefficient of variation (CV) for gene expression within single cells across all genes (left) and for single genes across all cells (right) in both the true distribution (green) and posterior predictive simulations. X-axes limits are set to include all CVs from the true distribution and scHPF, and as many CVs from other methods as possible. Accompanying bar graphs show the maximum distances between the cumulative distributions of the true and simulated CV values, (the Kolomogorov-Smirnov (KS) statistic, lower is better). **(c)** Same as (b), but clipping impossible negative posterior predictive samples to zero.

In bulk RNA-seq, modeling over-dispersed gene expression data has proven essential to downstream analysis [17]. In scRNA-seq, expression data are over-dispersed both across genes in individual cells and for individual genes across cells. We evaluated how well different factorization methods captured single cell expression variability using a posterior predictive check (PPC). PPCs provide insight into a generative model’s goodness-of-fit by comparing the observed dataset to simulated data generated from the model. More formally, PPCs sample simulated replicate datasets X_rep_ from a generative model’s posterior predictive distribution and use a modeler-defined test statistic to evaluate discrepancies between X_rep_ and the true data, X_obs_ [24]. For each dataset, normalization, and generative factorization method (scHPF, PCA, FA and ZIFA), we sampled ten replicate expression vectors per cell. After converting samples from models on normalized data back to molecular counts (Methods), we computed the coefficient of variation (CV) for all genes in each cell and each gene across all cells. Finally, we averaged each cell and gene’s CVs across the ten replicate simulations. In all three datasets, scHPF more closely matched the observed data’s variability than other methods (**Fig. 2b**, **Fig. S3**). We noticed that many samples from PCA and FA had physically impossible negative values. When we corrected these values by clipping them to zero, PCA and FA’s estimates of variability across cells collapsed toward zero (**Fig. 2c**). This collapse suggests that PCA and FA’s ability to model over-dispersion in scRNA-seq data depends on placing probability mass on negative gene expression levels.

### Application to Spatially Sampled scRNA-seq from High-Grade Glioma

As a demonstration, we applied scHPF to 6,109 single cell expression profiles from the core and invasive edge of a high-grade glioma. High-grade gliomas (HGGs), the most common and lethal brain malignancies in adults [25], are highly heterogeneous tumors with complex microenvironments. In HGG, malignant cells invade the surrounding brain tissue, forming diffusely infiltrated margins that are impossible to fully remove surgically [26]. Although malignant cells in margins seed tumor recurrence and are the targets of post-operative therapy, most molecular characterization has focused on HGG cores. To investigate the transcriptional differences between cells in glioma’s core and margins, we used an MRI-guided procurement technique [26] and scRNA-seq to profile 3,109 cells from an HGG core and 3,000 cells from its margin. While recent efforts are beginning to shed light on the differential expression between glioma’s core and margins [26, 27], few studies involve this kind of spatial sampling.

Glioma cells typically resemble glia at the level of gene expression, and our prior work characterizing HGGs with scRNA-seq revealed co-occurring malignant subpopulations resembling astrocytes, oligodendrocyte progenitors (OPCs), and neuroblasts [28]. Consistent with these findings, clustering and aneuploidy analysis (Methods, **Fig. S4-5**) revealed malignant subpopulations that expressed markers of astrocytes, OPCs, neuroblasts, and dividing cells as well as nonmalignant populations of myeloid cells, oligodendrocytes, endothelial cells, and pericytes (**Fig. 3a-b**, **S4-5**). In the spatially resolved samples, malignant subpopulations had dramatically different abundances across regions (Figure S5h). Astrocyte-like glioma cells were over two-fold more abundant in the margin biopsy, while OPC-like and cycling populations were nearly three and four-fold better represented in the core biopsy. All seventeen neuroblast-like glioma cells localized to the tumor core.

**Figure 3:**
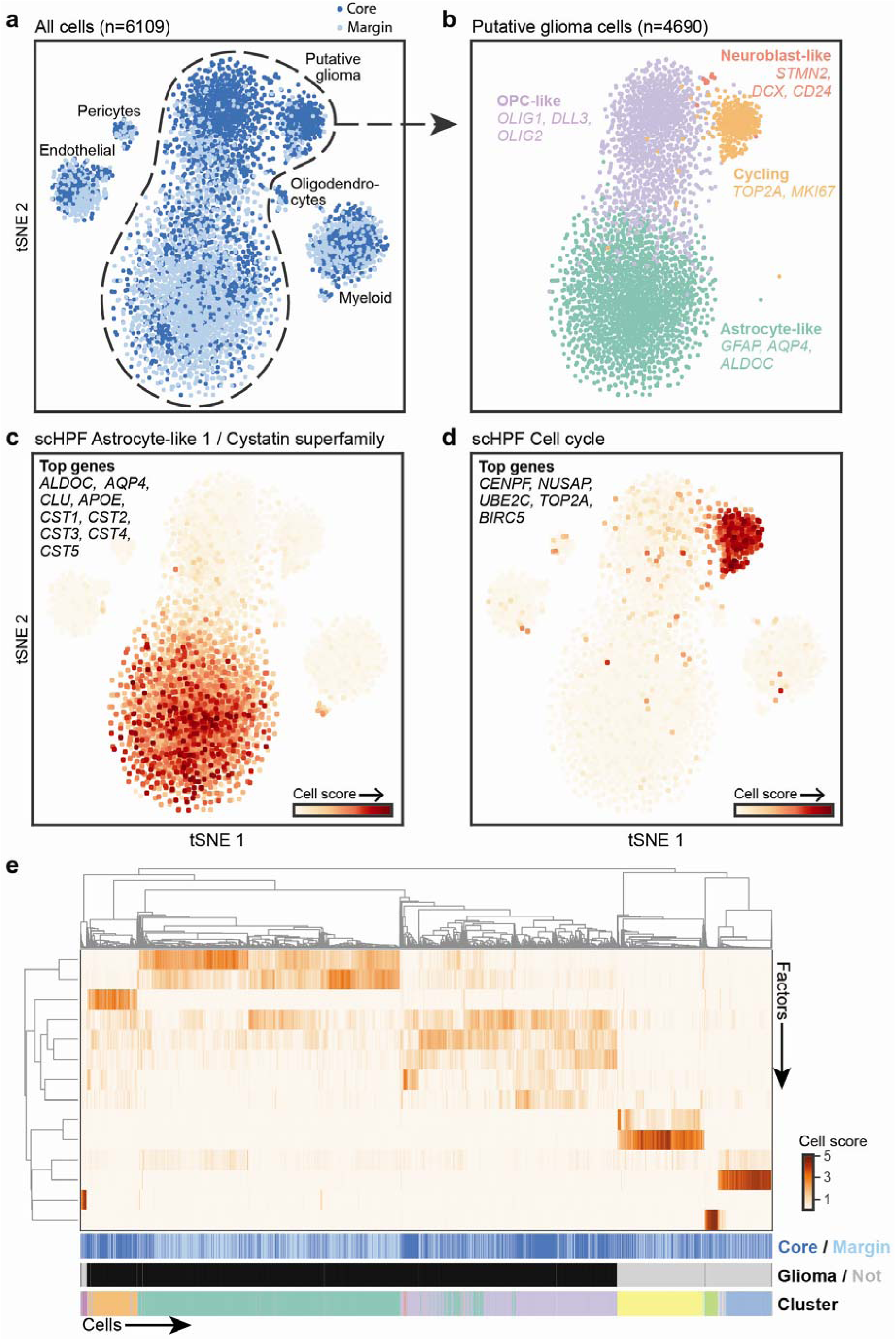
scHPF agrees with conventional analysis for regionally identified scRNA-seq of a High-Grade Glioma (HGG). **(a)** t-distributed Stochastic Neighbor Embedding (tSNE) [42]plot of cells from the core (navy) and margin (light blue) of an HGG reveal both malignant and non-malignant subpopulations (Methods). Labels were determined using malignancy score, clustering, and differential expression (Figures S5-6, Methods). **(b)** tSNE representation of putative glioma cells colored by cluster highlights astrocyte-like, OPC-like, neuroblast-like, and cycling subpopulations. **(c)** tSNE representation of all tumor cells colored by scHPF cell scores for one of two astrocyte-like factors. Nine out of the top 30 highest scoring genes are highlighted. **(d)** Same as (c), but for a cell cycle factor identified by scHPF. The five top-scoring genes in the factor are listed. **(e)** Main heatmap shows hierarchical clustering of cells’ scores for each factor. Top colorbar indicates the cell’s region: core (navy) or invasive edge (light blue). Second colorbar shows putative neoplastic status. Bottom colorbar indicates cluster.

Applied to the same dataset, scHPF identified at least one factor associated with every cell type, as well as physiological processes like translation, cell cycle and stress response (**Fig. 3c-d**, **S6b-d**). Hierarchical clustering of cells’ scores across factors recapitulated both Louvain clustering and malignant status (**Fig. 3e**), and factors associated with malignant subpopulations had regional biases across glioma cells that were consistent with glioma subpopulations’ differential abundance across regions (**Fig. S7a**). Therefore, scHPF captures major features identified by standard analyses for this dataset.

Some scHPF factors’ scores varied within the subpopulations identified by clustering. For example, two myeloid-associated factors that ranked pro-inflammatory cytokines and S100-familly genes highly (**Fig. S6a**), respectively, were correlated across all cells (r=0.66, p<10^−100^) but anticorrelated within the myeloid cluster (r=−0.59, p<10^−71^). Together, they appeared to represent a continuum of immune activation (**Fig. 4a-c**). This phenotypic gradient within an individual tumor is reminiscent of the variable myeloid states observed across different patients in previous studies of glioma [28-31].

**Figure 4:**
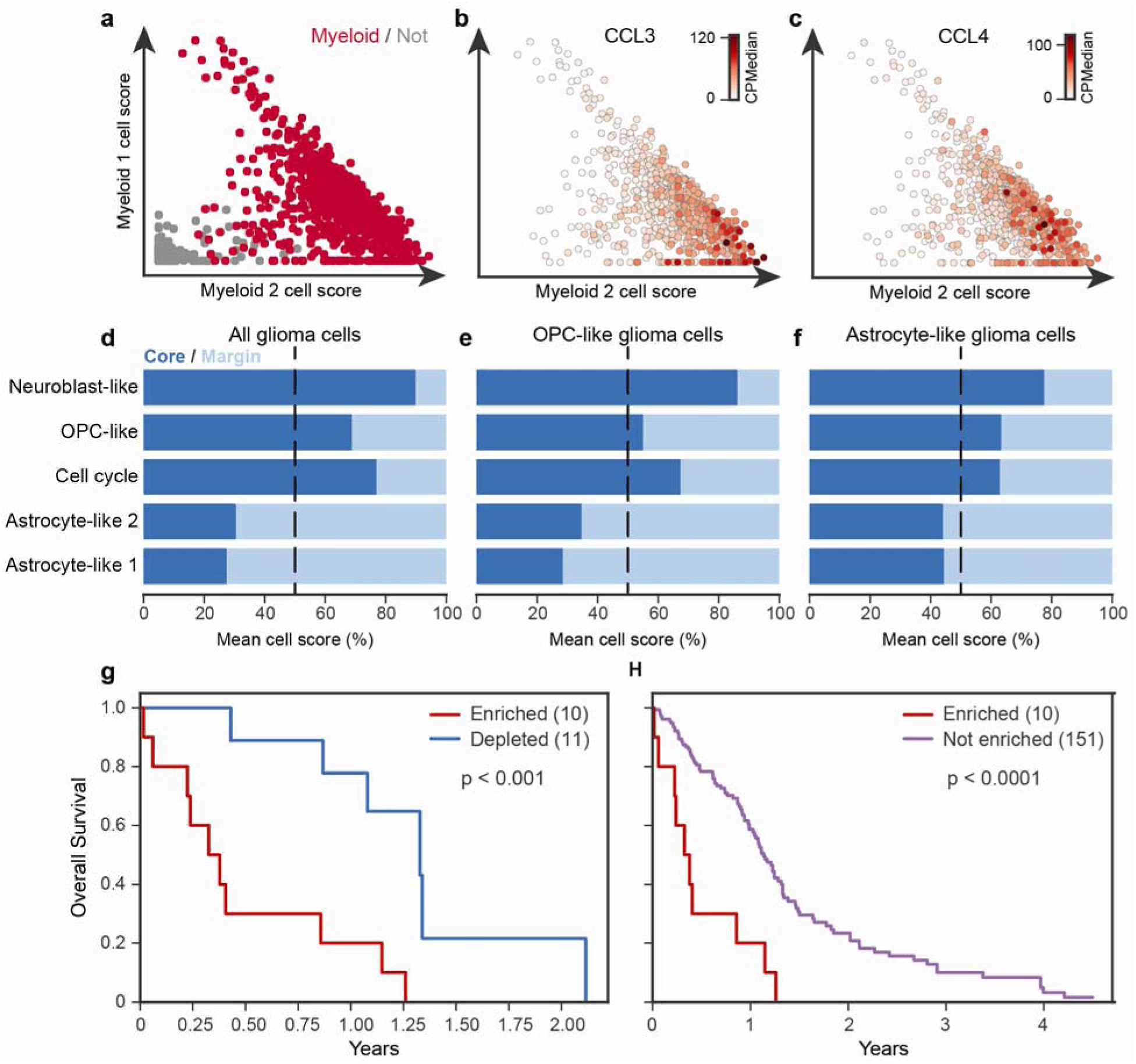
scHPF identifies finely resolved and novel, regionally associated features of HGG. **(a)** Scores for myeloid factor 1 (y-axis) vs. myeloid factor 2 (x-axis) for cells in the myeloid Louvain cluster (crimson) and all other cells (gray). Expression of proinflammatory cytokines CCL3 **(b)** and CCL4 **(c)** for cells in the myeloid subpopulation show a gradient of activation. **(d-f)** Factor score bias between the core (navy) and margin (light blue) in all glioma cells (d), OPC-like glioma cells (e), and astrocyte-like glioma cells (f). Mean cells scores in each region are scaled to sum to 100. Biases are driven by the top genes in each factor (Fig. S7d-f). **(g & h)** Kaplan-Meir curves show survival differences in TCGA for donors enriched (red), not enriched (purple), and depleted (blue) for the 25 top scoring genes in astrocyte-like factor 1 (Methods).

While scHPF factors had regional biases that reflect overall compositional differences between the core and margin biopsies, glioma cells’ scHPF factor scores also exhibited regional biases *within* the malignant subpopulations defined by clustering (**Fig. 4d-f**, **S7a**). For example, OPC-like glioma cells in the tumor core had significantly higher scores for the neuroblast-like, OPC-like, and cell cycle factors than their counterparts in the margin (Bonferroni corrected p<10^−84^, p<10^−6^ and p<10^−6^ respectively by the Mann-Whitney U test), whereas OPC-like glioma cells in the margin had higher scores for the two astrocyte-like factors (p<10^−49^ and p<10^−69^ for astrocyte-like factors 2 and 1, respectively). These differences are driven by the highest scoring genes in each factor (**Fig. S7b**), and astrocyte-like glioma cells followed a similar pattern. An alternative method of determining cellular subpopulations, where cells were assigned to the subpopulation with which their highest scoring factor was associated, also preserved the regional biases (**Fig. S7c**). This analysis suggests that, in this case, cells in the same malignant subpopulations but different tumor regions may have subtly different lineage resemblances.

As cells from the HGG margin remain after surgery and seed aggressive recurrent tumors, we investigated whether regionally-biased transcriptional signatures derived from scHPF factors were associated with survival in The Cancer Genome Atlas (TCGA) [32]. Restricting the analysis to glioblastoma (GBM), we identified patients enriched and depleted for the top genes in each factor (Methods). Survival analysis revealed significantly shorter overall survival (1 year median difference) for patients enriched for a margin-biased scHPF astrocyte-like signature (**Fig. 4g,h**), which included astrocytic markers *ALDOC*, *CLU* and *SPARCL1* [33-35], as well as cystatin super-family members *CST1* though *CST5* (**Fig. 3c**, **S6a**). Cystatin C (*CST3*) is highly expressed in mature human astrocytes [33, 35] and is induced in Alzheimer’s disease and epilepsy [36-38], raising the possibility that astrocyte-like glioma cells may be responding to the same cues or stresses that reactive astrocytes encounter in these disorders. Although it is difficult to determine which cells are responsible for an expression signature in bulk RNA-seq data, top scHPF astrocyte-like factor 1 genes were better correlated with molecular markers of tumor cells than other cells in the tumor microenvironment (**Fig. S8**), suggesting that glioma cells express those genes.

## Discussion

Conventional approaches to analyzing scRNA-seq data use predefined structures like clusters or pseudo-temporal orderings to identify discrete transcriptional programs associated with particular subpopulations and pseudo-temporally coupled gene signatures. However, gene expression programs may vary independently of these structures across complex populations. scHPF complements conventional approaches, allowing for *de novo* identification of transcriptional programs directly from a matrix of molecular counts in a single pass. By explicitly modeling variable sparsity in scRNA-seq data and avoiding prior normalization, scHPF achieves better predictive performance than other *de novo* matrix factorization methods while also better capturing scRNA-seq data’s characteristic variability.

In scRNA-seq of biopsies from the core and margin of a high-grade glioma, scHPF recapitulated and expanded upon molecular features identified by standard analyses, including expression signatures associated with all of the major subpopulations and cell types identified by clustering. Importantly, some lineage-associated factors identified by scHPF varied within or across clustering-defined populations, revealing features that were not apparent from cluster-based analysis alone. Clustering analysis showed that astrocyte-like glioma cells were more numerous in the tumor margin while OPC-like, neuroblast-like, and cycling glioma cells were more abundant in the tumor core. scHPF not only recapitulated this finding, but also illuminated regional differences in lineage-resemblance within glioma subpopulations. In particular, both OPC-like and astrocyte-like glioma cells in the tumor core had a slightly more neuroblast-like phenotype than their more astrocyte-like counterparts in the margin. Finally, we discovered a margin-biased gene signature enriched among astrocyte-like glioma cells that is highly deleterious to survival in GBM.

Massively parallel scRNA-seq of complex tissues in normal, developmental, and disease contexts has challenged our notion of “cell type” [39], particularly as highly scalable methods provide ever-increasing resolution. Further, gene expression programs essential to tissue function may be highly cell type-specific or might vary continuously within or across multiple cell types. Conventional graph- and clustering-based methods provide invaluable insight into the structure of complex cellular populations, and much can be learned from projecting single-cell expression profiles onto these structures. scHFP effectively models the nuanced features of scRNA-seq data while identifying highly variable gene signatures, unconstrained by predefined structures such as clusters or trajectories. We anticipate that scHFP will be a complementary tool for dissecting the transcriptional underpinnings of cellular identity and state.

## Methods

### Single-cell Hierarchical Poisson Factorization

The generative process for single-cell Hierarchical Poisson Factorization, illustrated in **Figure S1a**, is:

1. For each cell *c:*

a. Sample capacity *ξ_c_* ~ *Gamma*(*a*′*, b*′)
b. For each factor *k*:
c. 
  i. Sample weight *θ_c,k_* ~ *Gamma*(*a, ξ*)
2. For each gene *g*:

a. Sample capacity *η_g_* ~ *Gamma*(*c*′*, d*′)
b. For each factor *k*:

a. Sample weight *β_g,k_* ~ *Gamma*(*c, η_g_*)
3. For each cell *c* and gene *g*, sample observed expression level

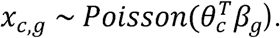

Where *x* is a discrete scRNA-seq expression matrix.

For *de novo* gene signature identification, we define each cell *c*’s score for each factor *k* as

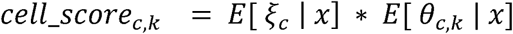

and each gene *g*’s score for each factor *k* as

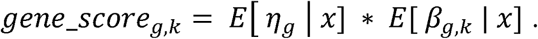

This adjusts factor loadings for the learned transcriptional output of their corresponding cell or gene. Finally, we rank the genes in each factor by their scores to identify *de novo* patterns of coordinated gene expression (e.g. **Fig S6a**). Cell’s scores, for example those plotted **Figures 3c-d** and **S6b-d**, indicate a cell’s association with the factor.

### Inference

We use Coordinate Ascent Variational Inference to approximate *p*(*ξ*,*θ*,*η*,*β* | *x*), the posterior probability of the model parameters given the data [1]. Hyperparameters *a*’, *b*’, *c*’ and *d*’ are set to preserve the empirical variance to mean ratio of the total molecules per cell or gene in the Gamma distributions from which *ξ* and *η* are drawn. Specifically, we set

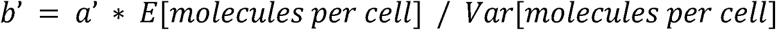

and

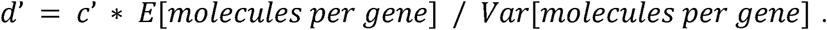

To preserve sparsity, we fix *a* and *c* to 0.3 and *a’* and *c*’ to 1. In this scheme, we find the algorithm largely insensitive to small changes in the hyperparameters. We initialize the variational distributions for *ξ*, *θ*, *η*, *β* to their priors times a random multiplier between 0.5 and 1.5.

### Selection of number of factors

In actually usage, such as the for the high-grade glioma demonstration in this paper, we select the number of factors *K* such that (1) the model’s log likelihood has converged (**Fig. S9a**) and (2) each well-defined cell-type in the dataset is most strongly associated with at least one factor with which no other cell-type is most strongly associated (**Fig. S9b-d**). For benchmarking experiments, to avoid biasing results toward any one method, we set the number for factors to the smallest multiple of five greater than the number of clusters for the PBMC and Matcovitch *et al*. datasets, and to five for TS543 (**Table S1**). However, predictive performance was robust to a range of values for *K* (**Fig. S2**).

### Benchmarking

Log-normalization was applied by adding 1 to molecular counts and then taking the logarithm. Counts per median (rate-normalization) were calculated by normalizing the molecular counts in each cell to sum to 1 and then multiplying all values by the median number of molecules per cell. For log-rate-normalization, we performed the log-normalization procedure described above on rate-normalized data. PCA, NMF, and FA were applied using the scikit-learn python package, with default parameters [40]. To test ZIFA, we used its authors’ block_ZIFA implementation with parameter p0_thresh=1 and otherwise default settings. Prior to training, we randomly selected 4% of nonzero expression values to use as a held-out test set and 2% as a validation set. The remaining data were used as a training set. By holding out only a small portion of data, we aimed to minimally impact datasets’ native sparsity structure. As these test and validation sets were small compared to the training set, we evaluated methods’ predictive performance on at least three randomly chosen partitions of the data into training, validation and test sets. We ran each method-normalization pair with ten random initializations on each training set and selected the run with the lowest mean absolute error on the corresponding validation set. Due to ZIFA’s long runtime (~23 hours per initialization on TS543), we only ran it with five initializations per training set and for only one value of *K* (**Fig. S2**).

We generated posterior predictive samples from scHPF by sampling latent representations *θ _c_* and *β _g_* from the variational posterior and taking the inner product. For PCA, FA, and ZIFA, we sampled latent representations and expression values according to their underlying generative models [41]. For each method, normalization, and dataset, we sampled ten M x N datasets. Samples from models on normalized data were inverse transformed back to molecular counts before calculating column and row coefficients of variation. For example, samples from PCA on log-normalized data were added to −1 and then exponentiated before calculating coefficients of variation. Each gene and cell’s coefficient of variation was averaged across ten replicate posterior predictive simulations. The Kolomgorov-Smirniov test statistic was calculated using the python package scipy.

### Published scRNA-seq datasets

The filtered PBMC dataset, using Chromium v2 chemistry, was downloaded from 10x Genomics (https://support.10xgenomics.com/single-cell-gene-expression/datasets/2.1.0/pbmc4k). Molecular counts for Matcovitch *et al*. were retrieved from GEO ascension GSE7918.

### Preparation of TS543 glioma neurospheres

TS543 cells were plated at density 1 x 104 viable cells/cm^2^ and grown as neurospheres with NeuroCult™ NS-A Basal Medium supplemented with NeuroCult™ NS-A Proliferation Supplement, 20ng/ml EGF, 10ng/ml bFGF, and 0.0002% Heparin (Stem Cell Technologies). When diameters of neurospheres reached to approximately 100μm, neurospheres were dissociated to single cells with mechanical force by pipetting 30-50 times.

### Radiographically-guided biopsies of high-grade glioma

Human glioma surgical specimens were procured from de-identified patients through a protocol approved by the Columbia Institutional Review Board (IRB). Radiographically-guided biopsies were obtained as described in [26]. Briefly, the patient studied here presented with FLAIR hyper-intense, non-contrast-enhancing tissue along the surgical trajectory based on MRI between the craniotomy site and gadolinium contrast-enhancing border of the lesion. This region was biopsied and comprised the tumor margin specimen described above. A region of the contrast-enhancing core of the lesion was also biopsied and comprised the tumor core specimen.

### Whole-genome sequencing

Low-pass whole genome sequencing (WGS) was conducted as described in [28]. Briefly, we homogenized tissue with a Dounce and extracted DNA and RNA with a ZR-Duet Kit (Zymo) according to the manufacturer’s instructions. For the normal control, DNA and RNA were extracted using the same kit from peripheral blood mononuclear cells. WGS libraries were constructed by *in vitro* transposition using the Illumina Nextera XT kit and sequenced on an Illumina NextSeq 500 with 2×75 base paired-end reads to a depth of ~1x. Reads were aligned to the hg19 build of the human genome using *bwa-mem* and the coverage for each chromosome was quantified using *bedtools* after collapsing PCR duplicates with *samtools*. To generate the bulk WGS heatmap in **Fig. S5e**, we took the divided the normalized coverage of each chromosome in the tumor sample by that of the normal sample, normalized the resulting ratio by the median ratio across all chromosomes, and multiplied by two to estimate average copy number of each chromosome in the tumor sample.

### scRNA-seq data preprocessing

Reads for TS543 and HGG samples were processed into molecular count matrices as described in Yuan *et al*. [28]. For all benchmarking and scHPF analyses, we only considered protein-coding genes that were expressed in at least 0.1% of cells in the dataset, rounded to the next largest multiple of 5 (**Table S1**).

### Identification of malignant glioma cells

We identified malignantly transformed cells by two orthogonal methods. First, we clustered cells’ scRNA-seq profiles (see *Clustering and visualization)* and defined putative malignant cells using the genes most specific to each cluster (**Figure S4**, **S5a**). Next, we performed PCA of cells’ whole-chromosome expression and found that the first principal component, which we call the malignancy score, separated putatively transformed cells from non-malignant cells (**Fig. S5b-d**). For further validation, we computed putative glioma cells’ average chromosomal expression profiles relative to putative non-malignant cells and found that they were in good agreement with aneuploidies identified by low-coverage whole genome sequencing of bulk tissue from the tumor core (**Fig. S5e**).

### Clustering and visualization

Clustering, visualization and identification of cluster-specific genes was performed similarly to Yuan *et al*. [28], with an updated method for selecting genes detected in fewer cells than expected given their apparent expression level (likely markers of cellular subpopulations). Briefly, for variable gene selection only, we normalized the molecular counts for each cell to sum to 1. Genes were then ordered by their normalized expression values. For each gene *g* we calculated *f_g_*, the fraction of cells in the dataset that expressed *g*, and 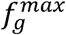, the maximum *f_g_* in a rolling window of 25 genes centered on *g*. 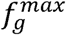 approximates the fraction of cells in which we expect to observe transcripts given *g*’s overall expression in the dataset. The scaled difference between *f_g_* and 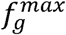 defines *g*’s dropout score:

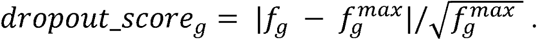

We selected marker genes with dropout scores that are either greater than 0.15 or at least six standard deviations above the mean, inclusively.

To cluster and visualize the data, we computed a cell by cell Spearman’s correlation matrix using the marker genes identified above. Using this matrix, we constructed a k-nearest neighbors graph (k=20), which we then used as input to Louvain clustering with Phenograph [4]. After clustering, we identified genes most specific to each cluster using a binomial test [5]. The same similarity matrix, transformed into a distance matrix by subtracting its values from 1, was used as input to tSNE for visualization.

### Regional biases

p-values for both factors and top scoring genes in each factor were calculated using the Mann-Whitney U-test and Bonferroni corrected for the total number of factors.

### Survival analysis

TCGA data for glioblastoma was downloaded from GDAC Firehose. Normalized expression values were log2(RSEM+ 1) transformed and each factor’s expression program was defined as its 25 highest scoring genes. We then calculated each program’s mean relative expression for each donor, and z-scored these values across donors. For each program, donors with z-scores greater than 1.5 were considered enriched, and all others were defined as not enriched. Patients with z-scores less than −1.5 were considered depleted. Kaplan-Meier curves and log-rank test p-values were generated with the Lifelines v0.11.1 Python module.

## Acknowledgements

We thank the Sulzberger Columbia Genome Center for assistance and resources for high-throughput sequencing. P.A.S. was supported by NIH/NCI grant R33CA202827, NIH/NIBIB grant K01EB016071, NIH/NCI grant U54CA209997, and a Human Cell Atlas Pilot Project grant from the Chan Zuckerberg Initiative. P.A.S., A.I., and A.L. are supported by NIH/NCI grant U54CA193313. P.A.S., P.C., and J.N.B. are supported by NIH/NINDS grant R01NS103473. DB is supported by ONR 133691-5102004, NIH 5100481-5500001084, and the John Simon Guggenheim Foundation. F.J.R.R. is supported by the EU Horizon 2020 programme (Marie Sklodowska-Curie Individual Fellowship, grant agreement 706760).

## Data availability

The single-cell RNA-Seq data generated for this study have been deposited in the Gene Expression Omnibus under accession GSE116621.

## Software Availability

Code is available at: https://github.com/simslab/scHPF

## Author contributions

HML and DMB conceived of the method. HML, DMB and PAS designed the study. JNB and PC procured glioma specimens. AL and AI prepared glioma samples. JY and YLC performed single cell sequencing. HML and FJRR wrote code. HML and PAS analyzed data. ECB performed whole-genome sequencing. HML and PAS wrote the manuscript with input from DMB, PC and FJRR. All authors read and approved the manuscript.

## Competing Interests

J.Y. and P.A.S. are listed as inventors on patent applications filed by Columbia University related to the microwell technology described here for single-cell RNA-Seq.

